# MYB transcription factor, NtMYB308, regulates anthocyanin and lignin content, and fungal tolerance in tobacco

**DOI:** 10.1101/2024.06.24.600478

**Authors:** Nivedita Singh, Shambhavi Dwivedi, Deeksha Singh, Pranshu Kumar Pathak, Prabodh Kumar Trivedi

**Affiliations:** Central Institute of Medicinal and Aromatic Plants (CSIR-CIMAP), Lucknow, India; Academy of Scientific and Innovative Research (AcSIR), Ghaziabad-201002, India

**Author notes:** **Corresponding author:** Prabodh Kumar Trivedi, CSIR-National Botanical Research Institute (CSIR-NBRI), Rana Pratap Marg, Lucknow-226001, INDIA; **; e-mail address:**. Responsible author for distribution of materials: Prabodh K. Trivedi. **Author contributions:** P.K.T. designed and supervised this study. N.S. designed and performed most of the experiments; SD, DS and PKP helped in conducting various experiments; P.K.T. and NS analyzed the data; N.S. and P.K.T. wrote the article; all authors read, contributed, and approved the article.

**Keywords:** AtMYB4-type, anthocyanin, biotic stress, gene expression, genome-editing, lignin, MYB308, negative regulation, tobacco

## Abstract

Anthocyanins are secondary metabolites synthesized through the phenylpropanoid pathway. They attract pollinators, possess antioxidant properties that scavenge free radicals during environmental stress, and provide protection against various stress conditions. Lignin, another secondary metabolite, plays crucial roles in providing mechanical support, facilitating water and solute transport, and protecting against pathogens. MYB transcription factors, particularly R2R3 MYBs, are key regulators of secondary metabolism, especially within the phenylpropanoid pathway. These factors act as both activators and repressors. The N-terminal region of R2R3-MYB repressors contains a conserved bHLH-binding domain, while the C-terminal domain is divergent and includes an EAR repressor domain. R2R3-MYB proteins notably target sequences such as the MYB-response element and AC elements. In this study, we identified and characterized the Nicotiana MYB transcription factor, NtMYB308, and explored its regulatory function in anthocyanin and lignin biosynthesis in tobacco. Our Virus Induced Gene Silencing (VIGS) and Protein-DNA interaction studies suggest that NtMYB308 is a negative regulator of anthocyanin and lignin biosynthesis by binding to the promoters of genes involved in these pathways. To validate our findings, we developed CRISPR/Cas9-based knockout mutant plants of tobacco, *NtMYB308^CR^*, which showed up-regulation of anthocyanin and lignin biosynthesis. Conversely, NtMYB308 overexpression (NtMYB308OX) plants exhibited the opposite effect. Enhanced anthocyanin and lignin levels in *NtMYB308^CR^* plants provided tolerance against the fungus *Alternaria solani*, while NtMYB308OX lines were susceptible. This study advances our understanding of the regulatory mechanisms governing anthocyanin and lignin biosynthesis and their role in biotic stress in tobacco.

**One Sentence Summary:** A R2R3 MYB transcription factor, NtMYB308, negatively regulates anthocyanin and lignin content, and fungal tolerance in tobacco.

## Introduction

Anthocyanins, synthesized through the phenylpropanoid pathway, play a crucial role in attracting pollinators and seed dispersers (Davies et al., 2012; Lev-Yadun et al., 2008). These compounds possess significant antioxidant properties, helping to scavenge free radicals and reactive oxygen species (ROS) during environmental stress, thereby offering protection under such conditions (Gould et al., 2004; Field et al., 2002; Gai et al., 2020). Moreover, anthocyanins are recognized for their health benefits as dietary supplements (Tsuda et al., 2012). The biosynthesis of anthocyanins is regulated by several key enzymes in the phenylpropanoid pathway, including phenylalanine ammonia-lyase (PAL), chalcone synthase (CHS), 4-coumarate-CoA ligase (4CL), flavonoid-3-hydroxylase (F3’H), dihydroflavonol-4-reductase (DFR), and anthocyanin synthase (ANS) (Sharma et al., 2020; Saito et al., 2013). In addition to anthocyanins, the phenylpropanoid pathway also synthesizes lignin, a complex polymer that forms a significant part of plant secondary walls. Lignin is essential for providing mechanical support, facilitating water and solute transport, and protecting against pathogens (Sharma et al., 2016; Mellerowicz et al., 2001; Boerjan et al., 2003). It is generated through the oxidative polymerization of three primary constituents: p-coumaryl, coniferyl, and sinapyl alcohols, collectively known as monolignols (Weng and Chapple, 2010). Recent functional analyses of transcription factors have enhanced our understanding of the intricate network of transcriptional regulation involved in the biosynthesis of flavonoids and secondary walls (Misra et al., 2010; Baucher et al., 2007; Demura & Fukuda, 2007; Zhong et al., 2008; Demura and Fukuda, 2007; Sharma et al., 2020; Gaddam et al., 2024).

Previous studies suggest that promoters of anthocyanin pathway genes, such as ANS and DFR, as well as lignin biosynthesis pathway genes like 4CL, CAD, and CCR, contain *cis*-regulatory elements. These elements include three AC motifs (AC-I: ACCTACC, AC-II: ACCAACC, and AC-III: ACCTAAC), which are binding sites for R2R3-MYB transcription factors that regulate these pathways (Borevitz et al., 2000; Jin et al., 2000; Tamagnone et al., 1998; Patzlaff et al., 2003). In plants, MYB transcription factors are pivotal regulators of secondary metabolism, particularly within the phenylpropanoid pathway, where they oversee the biosynthesis of anthocyanins and lignin (Pandey et al., 2014; Sharma et al., 2016; Bhatia et al., 2018; Cao et al., 2020). The synthesis of anthocyanins and proanthocyanidins (PAs) is intricately controlled by three categories of transcription factors i.e. R2R3-MYB factors, basic helix-loop-helix (bHLH) proteins, and WD40 repeat (WDR) proteins. These factors form the MBW complex, which is crucial for activating gene transcription (Koes et al., 2005; Ramsay and Glover, 2005; Naik et al., 2022). R2R3-The N-terminal region of these proteins contains a conserved bHLH-binding domain, facilitating interaction with bHLH proteins. The C-terminal domain, though more variable, features the characteristic C1 motif (LlsrGIDPxT/SHRxI/L) and C2 motif (pdLNLD/ELxiG/S), which are typical of subgroup 4 MYB proteins (Jin et al., 2000; Kagale and Rozwadowski, 2011). Additionally, the C-terminal domain includes the EAR repressor domain, further contributing to their regulatory functions (Naik et al., 2022). Apart from their transcriptional activation domains, MYB transcription factors also harbor regions for transcriptional inhibition, enabling them to act as repressors. The negative regulatory role of MYBs was initially demonstrated in snapdragon *Antirrhinum majus* (AmMYB308) (Tamagnone et al., 1998) followed by various other plant species.

R2R3-MYB repressors are divided into two main types: AtMYB4-like and FaMYB1-like (Yan et al., 2021). Examples of AtMYB4-type repressors include GbMYBF2, BrMYB4, AtMYB60, and AmMYB308, while FaMYB1-type repressors include TrMYB133, MYBC2, and PhMYB27. Apart from the C1 and C2 motifs, AtMYB4-like repressors feature additional motifs like the C3 or zinc-finger (ZnF) motif and the C4 motif, which is absent in FaMYB1-like repressors. FaMYB1-like repressors, such as PhMYB27, are known to participate in MBW complexes, whereas AtMYB4-like repressors directly bind to target gene promoters, as seen with MdMYB16 (Xu et al., 2017; Albert et al., 2014). This underscores the functional disparities and specific motifs associated with these R2R3-MYB repressor types. While some R2R3-MYBs facilitate lignin synthesis, Subgroup 4 members act as suppressors of genes involved in monolignol production (Jin et al., 2000; Legay et al., 2007; Fornale et al., 2006, 2010; Sonbol et al., 2009).

While MYBs have been identified as negative regulators of lignin biosynthesis in *Antirrhinum majus* and *Citrus sinensis* (Jia et al., 2018), their role in anthocyanin biosynthesis remains unstudied. In this study, we characterized NtMYB308 from tobacco and investigated its influence on both anthocyanin and lignin biosynthesis. Utilizing Virus-Induced Gene Silencing (VIGS), along with the development of overexpression and CRISPR/Cas9-based mutant plants, we examined the function of NtMYB308 in these pathways. Our findings suggest that NtMYB308 acts as an AtMYB4-type regulator, directly binding to the promoters of *NtANS, NtDFR, Nt4CL,* and *NtCAD*, thereby modulating both anthocyanin and lignin biosynthesis. Additionally, we observed that NtMYB308 contributes to fungal resistance in tobacco by regulating these pathways.

## Results

### Nicotiana MYB308 belongs to R2R3-MYB Subgroup 4 protein

The phenylpropanoid pathway is rigorously regulated by various protein families and transcription factors, ensuring specificity in secondary metabolite production (Albert et al., 2011; Xu et al., 2014; Naik et al., 2022). Both positive and negative regulators play crucial roles in the transcriptional control of secondary metabolite accumulation. This pathway synthesizes a wide array of metabolites, including flavonoids, anthocyanins, lignin precursors, phenolic acids, coumarins, and hydroxycinnamic acid esters (**Figure 1A**). While positive regulators for this pathway have been functionally characterized in several plants (Naik et al., 2022), there is limited information available on these factors in tobacco. The Nicotiana MYB repressor, NtMYB308, and its isoforms (NtMYB308-1, NtMYB308-2, and NtMYB308-3) were identified through the NCBI BLAST tool (https://blast.ncbi.nlm.nih.gov/Blast.cgi). A comparative alignment was performed with AtMYBL2 (Arabidopsis), PtrMYB182 (Poplar), EgMYB1 (Eucalyptus), and other R2R3-MYB repressors from various plants **(Supplemental Figure S1A)**. The protein sequence alignment revealed a highly conserved N-terminal R2R3 domain containing a bHLH-binding domain, while the C-terminal domain was significantly divergent. Two conserved motifs, the C1 motif (LlsrGIDPxT/SHRxI/L; Shen et al., 2012) and the C2 motif (pdLNLD/ELxiG/S), were identified as signature motifs for subgroup 4 MYB proteins due to their possession of an EAR repressor domain (Kagale and Rozwadowski, 2011; Jin et al., 2000) **(Supplemental Figure S1A)**. The Nicotiana MYB repressors share the LxLxL-type EAR motif with other phenylpropanoid MYBs involved in lignin/anthocyanin biosynthesis. Specifically, NtMYB308-1, NtMYB308-2, and NtMYB308-3 contain the LNLEL motif, similar to EgMYB1, TrMYB133, and PpMYB17. Like NtMYB308, many phenylpropanoid MYB repressors, such as AtMYB4 and EgMYB1 (Jin et al., 2000; Legay et al., 2010), contain C4 and zinc-finger motifs. However, these motifs are absent in repressors such as VvMYBC2-L1, PhMYB27, PtrMYB182, and FaMYB1.

**Figure 1.**
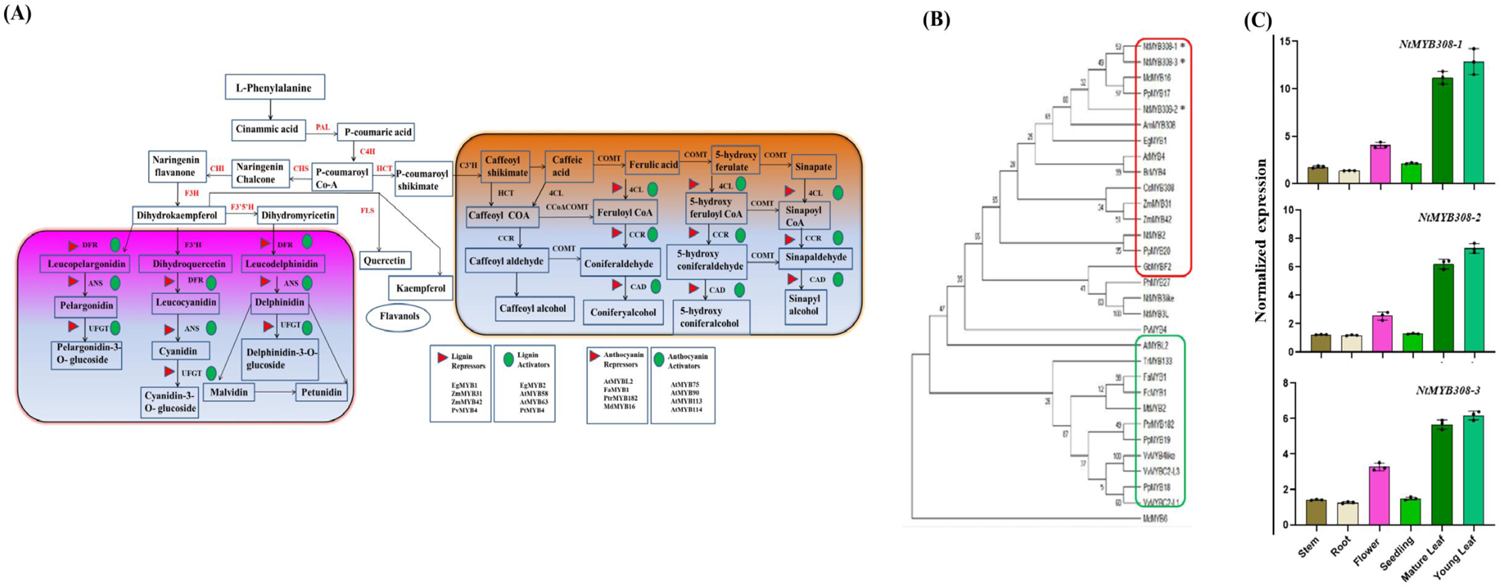
Phylogenetic analysis of NtMYB308 and tissue-specific expression. **A**, Phenylpropanoid pathway depicting structural genes and pathway enzymes. Enzymes in each step: PAL, phenylalanine ammonialyase; C4H, cinnamic acid 4-hydroxylase; 4CL, ρ-coumaroyl-CoA synthase; CHS, chalcone synthase; CHI, chalcone-flavanone isomerase; F3H, flavanone 3-hydroxylase; F3′H, flavanone 3′-hydroxylase; DFR, dihydroflavonol 4-reductase; ANS, anthocyanidin synthase; HCT, hydroxycinnamoyl-CoA:skimimate/quinatehydroxycinnamoyltransferase; C3’H, ρ-coumarate 3’-hydroxylase; COMT, caffeic acid 3-O-methyltransferase; CCR, cinnamoyl-CoA reductase; F5H, ferulate 5-hydroxylase; COMT, caffeic acid O-methyltransferase; CAD, cinnamyl alcohol dehydrogenase. ρ-Coumaroyl CoA is at the intersection point of both metabolic pathways i.e. anthocyanin synthesis or lignin synthesis. **B,** Phylogenetic tree of Nicotiana MYB308-1, MYB308-2, MYB308-3 and related functionally characterized R2R3-MYB transcription factors (sequence obtained from NCBI database) from across species of different families, constructed from the N-terminal DNA-binding domains using the maximum likelihood method. Stars indicate the Nicotiana repressor-like MYBs i.e. NtMYB308-1, NtMYB308-2 and NtMYB308-3 which falls in AtMYB4 clad. **C,** Tissue-specific expression of NtMYB308-1, NtMYB308-2 and NtMYB308-3. Normalized expression of NtMYB308 and its isoforms in different tissue (seedling, leaf, stem, root and flower). Actin was used as the endogenous control to normalize the relative expression levels. The statistical analysis was performed using two-tailed Student’s t-tests. Data are the means±SE of three biological (n=3) and three technical replicates. The error bars represent standard deviations. The asterisks indicate significant differences: *P<0.1; **P<0.01; ***P< 0.001.

A phylogenetic analysis was conducted on known MYB repressors involved in regulating anthocyanin and lignin biosynthesis to elucidate their relationship with NtMYB308. The analysis suggested two major clades, with Nicotiana MYB308 clustering in the clade containing AtMYB4-type R2R3-MYB factors. This clade includes MdMYB16, EgMYB1, AmMYB308, PpMYB17, PpMYB20, AtMYB4, CsMYB308, ZmMYB31, and ZmMYB42 (**Figure 1B**). In contrast, the other clade predominantly comprises FaMYB1-type R2R3-MYB factors, such as FaMYB1, FcMYB1, PhMYB27, VvMYBC2-L3, VvMYBC2-L1, and PtrMYB182. Therefore, the phylogenetic analysis indicates that NtMYB308 belongs to the AtMYB4-type subgroup 4 MYB repressor protein family. Expression analysis of NtMYB308 and its isoforms revealed similar patterns across various tissues. Constitutive expression was observed in leaves, flowers, stems, roots, and seedlings, with notably higher expression levels in leaves (**Figure 1C**). Interestingly, despite the general expectation of high anthocyanin accumulation in flowers and stems, the highest expression was found in leaves

### Silencing of NtMYB308 effects plant growth and enhances anthocyanin and lignin accumulation

The 3’ coding regions and untranslated regions (3’UTR) of three isoforms, NtMYB308-1 (211 bp), NtMYB308-2 (250 bp), and NtMYB308-3 (168 bp), were selected for VIGS analysis **(Supplemental Figure S1B)**. As a positive control, *NtPDS* (265 bp) was used. Upon infiltration of TRV:NtPDS into Nicotiana plants, a photobleached phenotype appeared in the systemic leaves approximately 22 days post-inoculation (d.p.i.) **(Supplemental Figure S2A)**. The silencing frequency was determined to be 62%, calculated by the number of plants infiltrated with TRV that exhibited the photobleached phenotype. Gene silencing success was further evaluated by counting the total number of leaves showing photobleached symptoms out of the total number of leaves on the plant at that stage, resulting in a 37% silencing efficiency. Notably, a significant down-regulation of *NtPDS* was observed in the photobleached leaf samples (30 d.p.i.), with silencing efficiency reaching approximately 80% compared to the control **(Supplemental Figure S2A-B)**. Plants infiltrated with the constructs targeting NtMYB308 exhibited characteristic viral symptoms (**Figure 2A**), and a notable increase in plant height compared to control plants **(Supplemental Figure S2C-D)**. Additionally, a significant alteration in total biomass was observed upon silencing of NtMYB308 **(Supplemental Figure S2E).** Expression analysis of infiltrated plants indicated a decrease in the accumulation of transcripts of all three isoforms of the NtMYB308 gene compared to control plants. Specifically, there was a significant reduction in transcript levels in plants infiltrated with TRV:NtMYB308-1, TRV:NtMYB308-2, and TRV:NtMYB308-3 by 74%, 65%, and 71%, respectively **(Supplemental Figure S2F)**.

**Figure 2.**
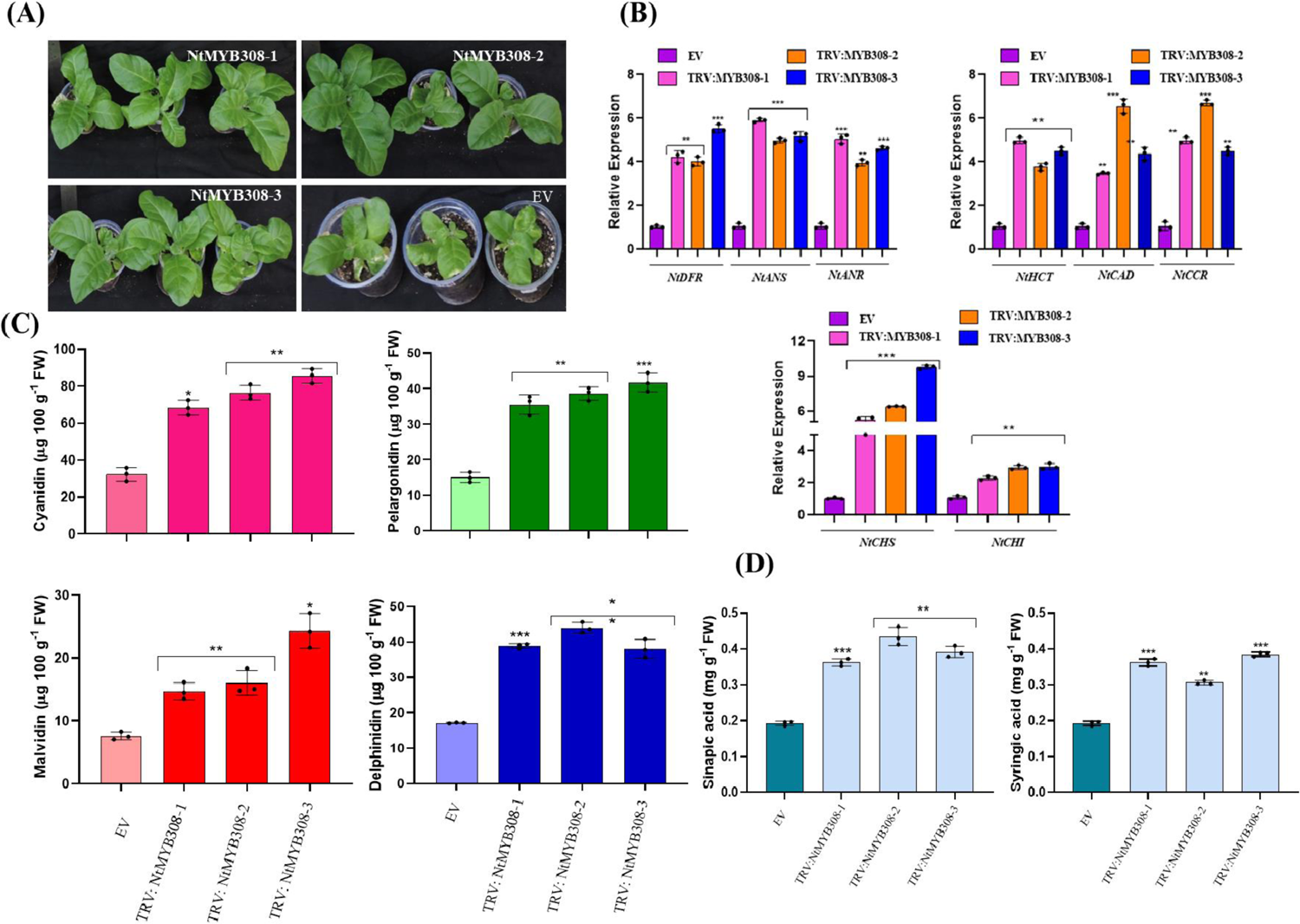
Tobacco Rattle Virus-mediated silencing of MYB308 in *Nicotiana tabacum*. **A,** Phenotype of the empty vector treated (Control) and TRV2:NtMYB308 treated *Nicotiana tabacum* plants at 30 d.p.i. The representative phenotypes of silenced plants using constructs with gene of interest. The phenotype exhibits viral infection, i.e. slight curling of leaves in silenced lines and empty vector (Control) at 30 d.p.i **B,** Relative expression of pheylpropanoid pathway genes (Upper panel: *NtDFR, NtANR, NtANS, NtHCT, NtCCR and NtCAD;* Lower panel: *NtCHS, NtCHI*). Actin was used as an internal control. **C,** Quantitative estimation of specific anthocyanin content (Delphinidin, Malvidin, Pelargonidin and Cyanidin) in TRV silenced lines of NtMYB308 and its isoforms and empty vector control. **D,** Quantitative estimation of phenolic content (syringic acid and sinapic acid) through HPLC in TRV silenced lines and empty vector control. Data are mean±SE of six biological (n=6) and three technical replicates. The error bars represent standard deviations. The asterisks indicate significant differences: *P<0.1; **P<0.01; ***P< 0.001.

The impact of down-regulating NtMYB308 on the expression of both early and late phenylpropanoid pathway genes was investigated in both control and TRV:NtMYB308-silenced plants. Transcript levels of early pathway genes, including *NtCHS* and *NtCHI*, as well as late pathway genes such as *NtANR, NtDFR, NtANS, NtCCR, NtHCT,* and *NtCAD*, exhibited significant increases in the silenced plants across all three NtMYB308 genes (NtMYB308-1, NtMYB308-2, and NtMYB308-3) (**Figure 2B**). Furthermore, the transcript levels of core anthocyanin pathway genes like *NtANS, NtANR, NtDFR*, and those involved in lignin biosynthesis such as *NtCCR, NtHCT,* and *NtCAD*, also showed notable increases upon silencing NtMYB308. These findings suggest that NtMYB308 acts as a negative regulator in both anthocyanin and lignin biosynthesis pathways. The silenced lines exhibited notably higher total anthocyanin content (**Supplemental Figure S2G)**. Analysis of specific anthocyanins, including delphinidin, malvidin, pelargonidin, and cyanidin, indicated a significantly enhanced accumulation of these compounds in NtMYB308-silenced plants relative to the control (**Figure 2C**). Furthermore, the assessment of total lignin content revealed enhanced accumulation of these compounds in the silenced plants compared to the control **(Supplemental Figure S2H)**. The infiltrated plants were analysed to detect the presence of the virus through infectivity analysis. Examination of the cDNA from infiltrated control and NtMYB308-silenced plants revealed the presence of coat protein-related amplicons, providing evidence of systemic viral infection **(Supplemental Figure S2I)**. These findings strongly suggest that the observed alterations in phenotype, transcript levels, and metabolite content are due to the silencing of isoforms of NtMYB308 in Nicotiana.

### Development and analysis of NtMYB308 overexpression and mutant plants

To investigate the influence of NtMYB308 on the regulation of the phenylpropanoid pathway, we developed overexpression lines (NtMYB308OX) and generated mutated plants (*NtMYB308^CR^*) through CRISPR/Cas9-based genome-editing approach. For the overexpression lines, we utilized the NtMYB308-1 gene, which was overexpressed under the control of the constitutive CaMV35S promoter. Among the T3 generation, three homozygous lines were identified for further analysis. Expression profiling revealed a significant increase in *NtMYB308* expression in transgenic lines compared to wild-type (WT), indicating successful overexpression (**Figure 3A**). To develop mutated plants (*NtMYB308^CR^*), we targeted the coding region common to all three isoforms (NtMYB308-1, NtMYB308-2, and NtMYB308-3). This allowed for the simultaneous mutation of all three isoforms using a single guide RNA, with separate screening conducted for each. Sequence analysis confirmed deletions in the coding regions of the NtMYB308 isoforms, resulting in frameshift mutations and truncated proteins. Remarkably, plants exhibited nucleotide deletions in all three isoforms of NtMYB308 **(Supplemental Figure S3A)**. The editing in nucleotide sequence and protein sequence are depicted in **Supplemental Figure S3A**. From the mutated lines, we selected those displaying mutations in all three isoforms, designated as *NtMYB308^CR-1,2,3^* (**Figure 3A**). Subsequently, homozygous *NtMYB308^CR-1,2,3^* plants devoid of Cas9 activity were identified and referred to *NtMYB308^CR^* in the manuscript further. Expression analysis demonstrated a significant reduction in transcript levels for all three isoforms in *NtMYB308^CR^*plants (**Figure 3A**). Moreover, we obtained edited plants with mutations specific to individual isoforms, resulting in decreased expression of the respective homolog **(Supplemental Figure S3B)**.

**Figure 3.**
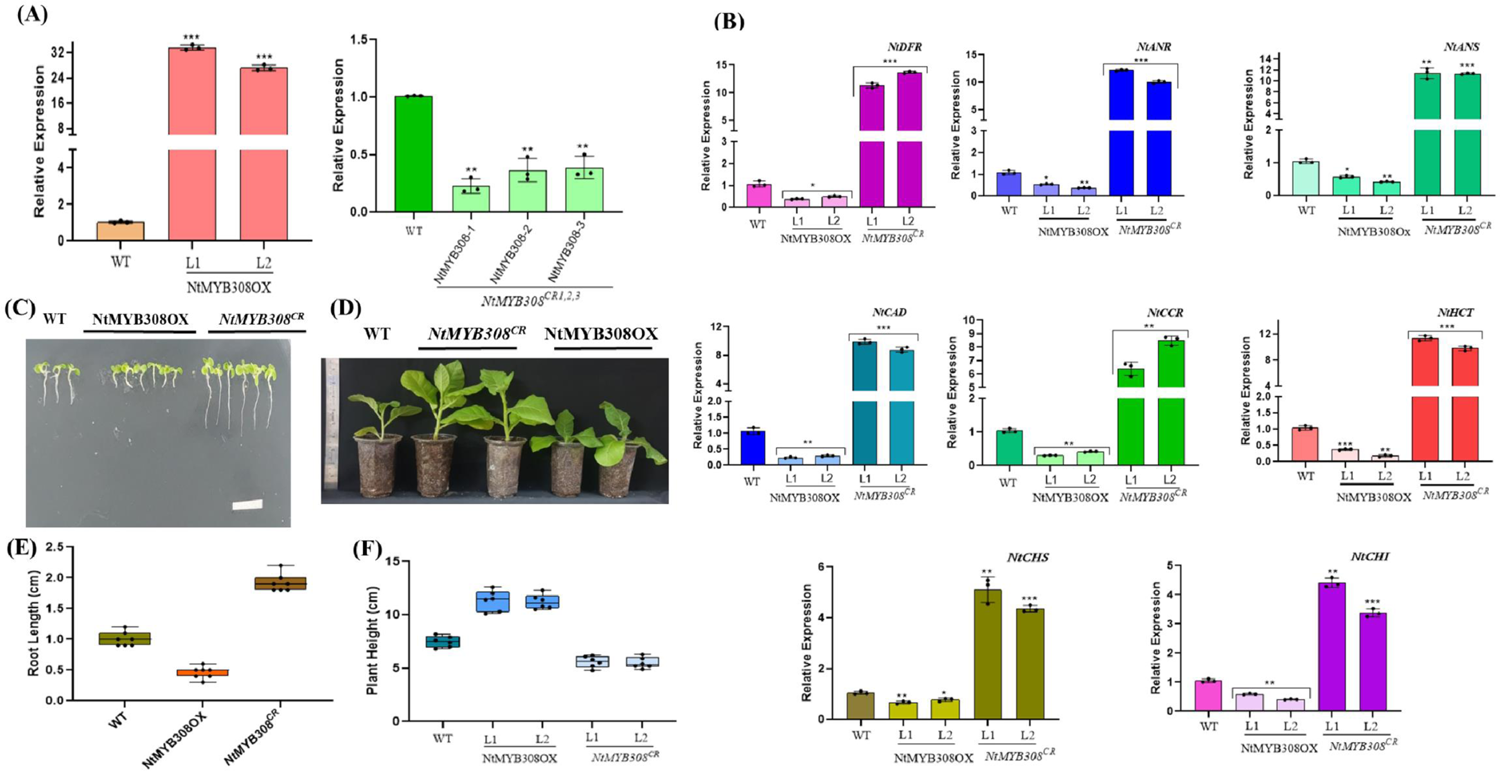
Expression of pathway genes in overexpression and mutant plants. **A**, Relative expression of NtMYB308-1 in NtMYB308OX and *NtMYB308^CR^* plants **B,** Relative expression of pheylpropanoid pathway genes (*NtCHS, NtCHI, NtDFR, NtANR, NtANS, NtHCT, NtCCR and NtCAD*) in WT, NtMYB308OX and *NtMYB308^CR^*lines. **C,** Phenotype of root length (cm) of 12 day old seedlings of WT, NtMYB308OX and *NtMYB308^CR^* lines (n=20 independent seedlings). **D,** Phenotype of plant height (cm) of 40 day old plants (n= independent plants). **E,** Root length (cm) of 12-day old seedlings of WT, NtMYB308OX and *NtMYB308^CR^* lines. **F,** Plant height (cm) of 40-day old plants (n= independent plants). Calculations of data were performed from three biological replicated independently per treatment with similar results. The statistical analysis was performed using two-tailed Student’s t-tests. The error bars represent standard deviations. The asterisks indicate significant differences: *P<0.1; **P<0.01; ***P< 0.001.

We conducted an expression analysis of key structural genes involved in anthocyanin and monolignol biosynthesis in both NtMYB308OX and *NtMYB308^CR^*plants. Notably, the expression levels of *NtANR, NtANS, NtDFR, NtCAD, NtCCR*, and *NtHCT* were significantly elevated in mutated plants, whereas they were reduced in overexpression lines compared to wild-type plants (**Figure 3B**). Similarly, the expression of early phenylpropanoid pathway genes, such as *NtCHS* and *NtCHI*, exhibited higher levels in mutant plants and decreased expression in overexpression lines (**Figure 3B**). Overexpression of NtMYB308 led to a downregulation of pathway gene expression, while its mutation resulted in the opposite effect, suggesting a regulatory role for NtMYB308 in modulating these metabolic pathways. This analysis indicates that NtMYB308 acts as a negative regulator of the phenylpropanoid pathway, encompassing anthocyanin and lignin biosynthesis.

### Modulation in NtMYB308 expression affects plant growth

Previous reports have demonstrated that overexpression of R2R3-MYB transcription factors, known as lignin repressors, can result in reduced growth and altered leaf morphology. In the case of NtMYB308OX transgenic plants, several phenotypic changes were evident, particularly in terms of root length and plant height. To further investigate the differences in root length between NtMYB308OX and *NtMYB308^CR^*plants, the root length of 12-day-old tobacco seedlings was measured. Remarkably, the root length of NtMYB308OX was significantly shorter compared to the wild-type, whereas the opposite trend was observed in *NtMYB308^CR^* plants (**Figure 3C** and **3E**). Notably, NtMYB308OX plants also exhibited a decrease in plant height, while *NtMYB308^CR^*plants displayed the opposite phenotype compared to wild-type plants. This observed reduction in phenotype in NtMYB308OX plants (**Figure 3D** and **3F**) mirrors findings reported for AtMYB4 and AmMYB308 in transgenic tobacco. Unlike transgenic tobacco overexpressing AmMYB308, we did not observe any white lesions on the leaves of our transgenic plants. In addition to alterations in plant height and root length, delayed flowering was evident in the transgenic lines, whereas genome-edited mutants exhibited early flowering compared to wild-type **(Supplemental Figure S3C)**. These findings collectively underscore the multifaceted role of NtMYB308 in regulating various aspects of plant growth and development.

### Reduced anthocyanin pigment in flower and mature plants overexpressing NtMYB308

Anthocyanin pigmentation exhibited a significant increase in *NtMYB308^CR^*plants, while it was notably reduced in NtMYB308OX plants compared to the wild-type (**Figure 4A**). To investigate anthocyanin accumulation, we examined 10-day-old tobacco seedlings, mature leaves from one-month-old plants, and flowers from various individuals. In flowers of *NtMYB308^CR^*plants, there was a pronounced abundance of total anthocyanin content and specific anthocyanins like cyanidin, delphinidin, malvidin and pelargonidin content **(Supplemental Figure S4A and Figure 4B**). Specific anthocyanins were found to be highly accumulated in leaves of mutants (**Figure 4C and Supplemental Figure S4B)** while a considerable reduction was observed in NtMYB308OX plants compared to wild-type counterparts. A similar pattern of anthocyanins was observed in seedlings **(Supplemental Figure S4D)**.

**Figure 4.**
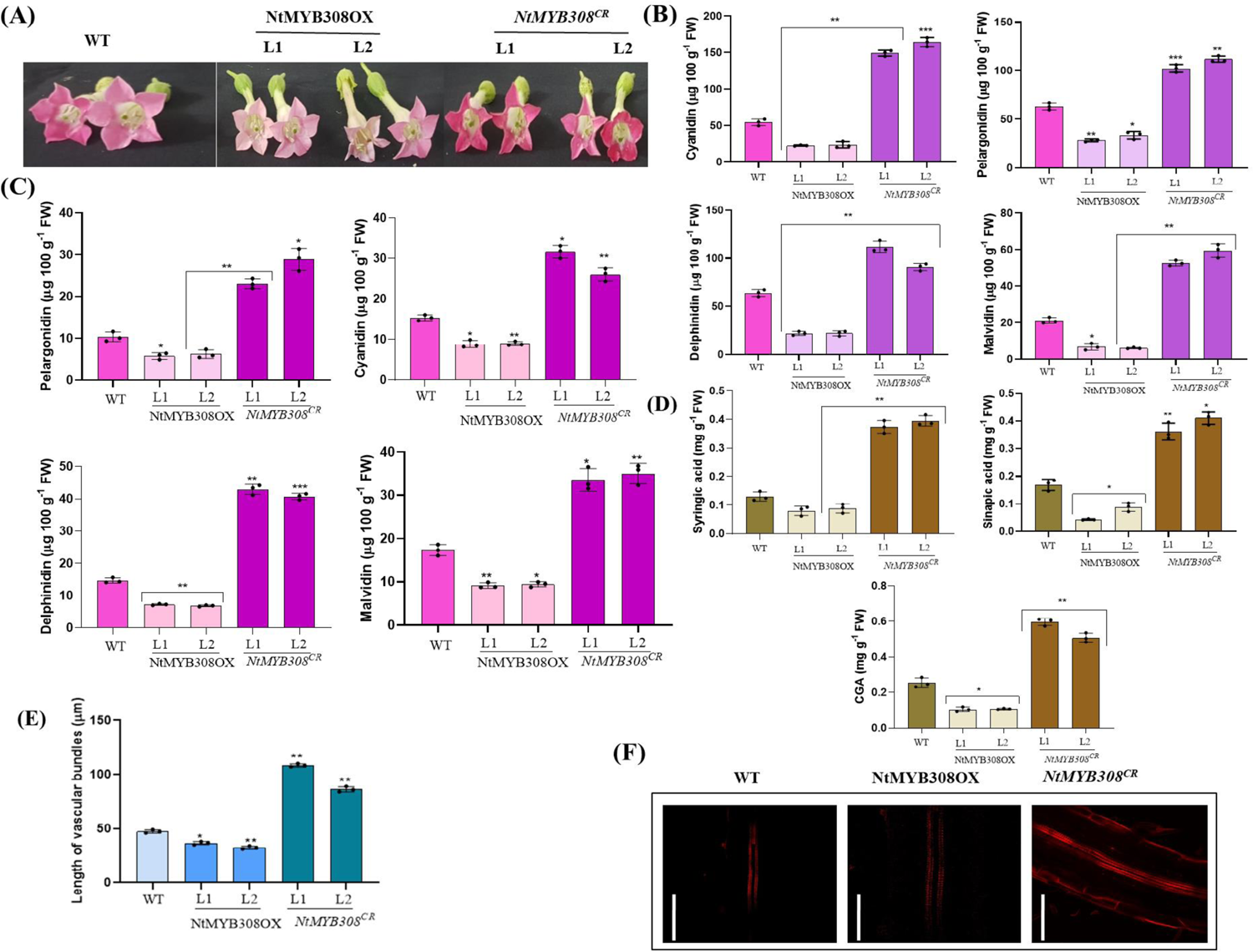
NtMYB308 inhibits the accumulation of anthocyanin content. **A**, Representative photograph of flower color alterations in petals of NtMYB308OX and*NtMYB308^CR^* lines as compared to WT. **B,** Quantification of specific anthocyanin content (Delphinidin, Malvidin, Pelargonidin and Cyanidin) in flowers of 40-day-old plant of WT, NtMYB308OX and *NtMYB308^CR^* plants. **C,** Quantification of specific anthocyanin content (Delphinidin, Malvidin, Pelargonidin and Cyanidin) in leaves of 40-day-old plant of WT, NtMYB308OX and *NtMYB308^CR^* lines. **D,** Quantification of specific phenolic content (Chlorogenic acid, Syringic acid and Sinapic acid) in stem of 45-day-old WT, NtMYB308OX and *NtMYB308^CR^* plants. **E,** Length of vascular bundles in 45-day-old stem of WT, NtMYB308OX and *NtMYB308^CR^* plants (*n* = 15) These experiments were repeated three times independently, with similar results. **F,** Primary roots of 20-day-old seedlings of WT, NtMYB308OX and *NtMYB308^CR^*lines were grown on half-strength Murashige and Skoog (MS) medium (Sigma) and stained with basic fuchsin for visualization of lignin and photographed at 20× magnification. Scale bar=50 μM. The statistical analysis was performed using two-tailed Student’s t-tests. Data are the means±SE of six biological (n=3) and three technical replicates. The error bars represent standard deviations. The asterisks indicate significant differences: *P<0.1; **P<0.01; ***P< 0.001.

### Modulation in lignin content and length of vascular bundles in NtMYB308 modulated plants

As NtMYB308 is a member of the MYB transcription factor family known for repressing the biosynthesis of lignin and other phenolic compounds, we analysed the levels of both total and specific lignin components. Among the most abundant soluble phenolic compounds and lignin polymers in tobacco stems are chlorogenic acid (CGA), sinapate esters, and ester-linked wall-bound syringyl aldehyde. To quantify total lignin content, stems from one-and-a-half-month-old NtMYB308OX and *NtMYB308^CR^* plants were analyzed, and results indicated higher lignin content in mutant plants as compared to WT and overexpression lines **(Supplemental Figure S4C)**. Similarly, upon overexpression of NtMYB308, there was a notable reduction in CGA, sinapic acid and syringic acid levels, whereas these compounds accumulated significantly in *NtMYB308^CR^* plants (**Figure 4D**) compared to wild-type plants. Moreover, examination of stem cross-sections from plants of the same age with lignin staining indicated reduced lignification in the interfascicular and vascular tissues of NtMYB308OX plants compared to wild-type, contrasting findings were observed in *NtMYB308^CR^* plants **(Supplemental Figure S4E)**. The length of vascular bundles was also investigated, revealing the highest length in mutants and the lowest in NtMYB308OX plants (**Figure 4E**). Given the pronounced inhibition of root growth observed in NtMYB308OX plants, we investigated whether the primary root growth inhibition induced by NtMYB308 was attributable to significant lignin deposition on the secondary cell wall. Visualization of lignin deposition in the region above the elongation zone of 20-day-old seedling roots stained with basic fuchsin revealed higher levels in *NtMYB308^CR^*plants and substantially lower levels in NtMYB308OX plants compared to wild-type (**Figure 4F**). These findings underscore the pivotal role of NtMYB308 in modulating lignin biosynthesis and its impact on plant growth and development.

### NtMYB308 interacts with promoters of genes involved in anthocyanin and lignin biosynthesis

To investigate the interaction between NtMYB308 and various promoters, we conducted Electrophoretic Mobility Shift Assays (EMSAs) using purified recombinant NtMYB308-6XHis-protein **(Supplemental Figure 5A-B**). Our focus was on evaluating whether NtMYB308 plays a role in the transcriptional regulation of lignin and anthocyanin biosynthesis genes by examining its interaction with AC elements and MBSIIG *cis*-elements present in the promoters of *Nt4CL, NtCAD, NtANS*, and *NtDFR*. *In silico* analysis of the promoters revealed the presence of AC elements upstream of the transcription start site. Specifically, we identified AC elements with core motifs (AC-I: ACCTACC, AC-II: ACCAACC, AC-III: ACCTAAC) within the 1.5-kb region upstream of the transcription start site of all the examined promoters **(Supplemental Figure S5C)**. Upon addition of the NtMYB308-His protein to probes derived from *Nt4CL, NtANS, NtDFR*, and *NtCAD*, a band shift was observed, indicating protein-DNA interaction. Conversely, no band shift was observed with mutated probes (m4CL, mCAD, mANS, mDFR), where the AC element had been disrupted **(Figure 5A-D)**. In the competition assay, as the concentration of unlabeled or cold probes increased from 10X to 50X, the intensity of binding diminished **(Supplemental Figure S5D)**. These results strongly suggest that NtMYB308 protein interacts with the promoters of *Nt4CL, NtCAD, NtDFR*, and *NtANS*, thereby regulating the expression of these genes and consequently influencing lignin and anthocyanin biosynthesis, respectively.

**Figure 5.**
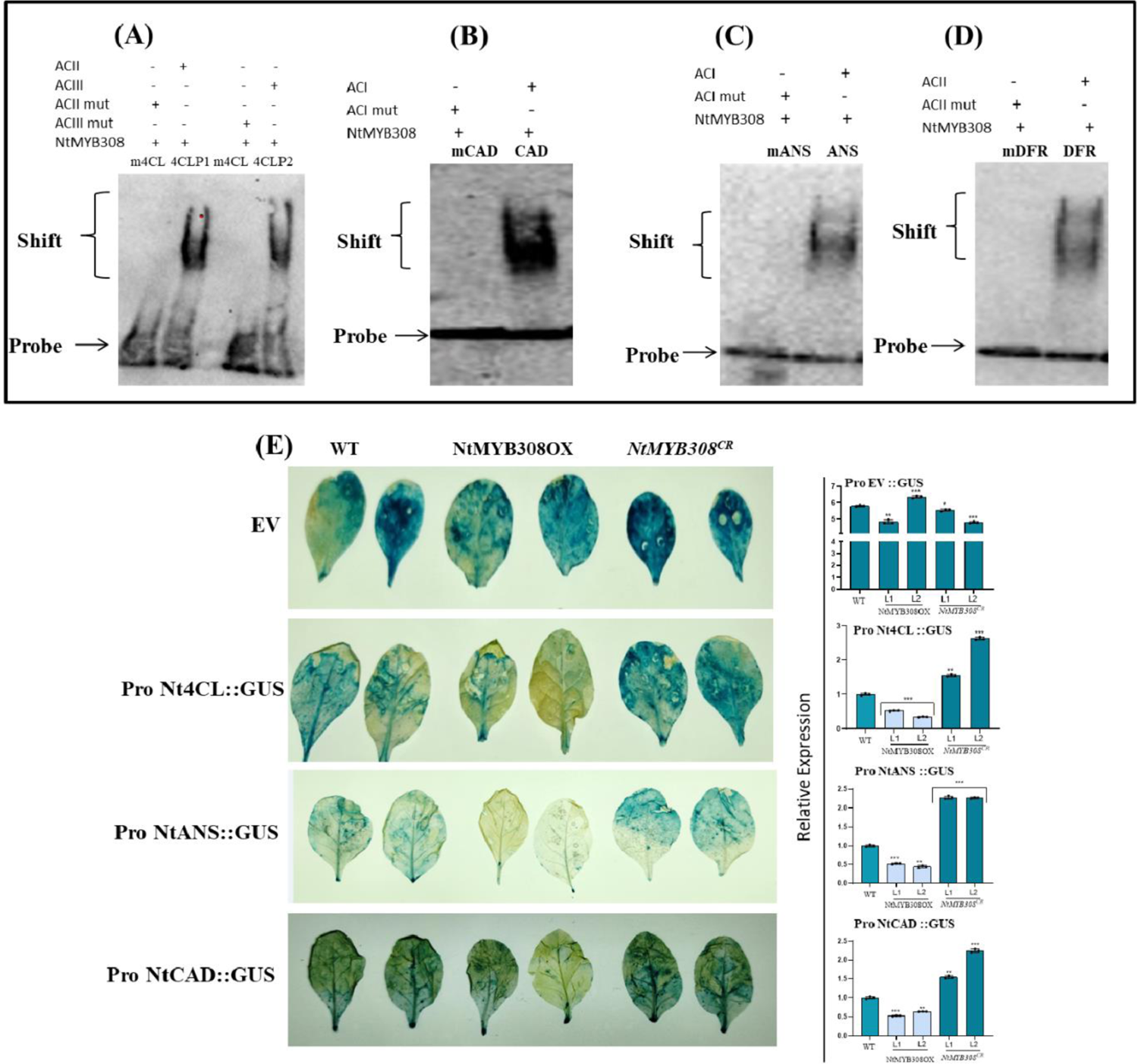
Interaction between NtMYB308 and promoters of *Nt4CL, NtANS, NtCAD* and *NtDFR* genes. **A-D**, EMSA (Electrophoretic Mobility Shift Assay) for the binding of NtMYB308-6X-His with core AC-DIG element (MYB308 binding site) present in the promoters of the *Nt4CL, NtCAD, NtANS* and *NtDFR* genes, respectively. Upper and lower arrows indicate shift and free probe respectively. **E,** Representative images of histochemical staining showing GUS (seen as blue color) and relative expression of GUS in leaves of WT, NtMYB308OX and *NtMYB308^CR^* lines of *Nicotiana tabacum* agroinfiltered with ProNt4CL::GUS, ProNtCAD::GUS, ProNtANS::GUS, and empty vector control. The experiment was repeated three times independently, with similar results. Statistical analysis was performed using two-tailed Student’s t-test. Error bars represent SE of means (n=3). Actin was used as endogenous control to normalize the relative expression levels. Error bars represent standard deviation. Asterisks indicate a significant difference, *P < 0.1, **P < 0.01, ***P < 0.001.

To elucidate the potential transcriptional regulation by NtMYB308 on target genes, NtMYB308OX, *NtMYB308^CR^*, and wild-type plants were agroinfiltrated with constructs carrying Nt4CLpro::GUS, NtCADpro::GUS, and NtANSpro::GUS, alongside an empty vector (pCAMBIA1303) as a control **(Supplemental Figure S6)**. Following agroinfiltration, GUS staining was scored and examined four days post-infiltration. Remarkably, GUS staining intensity increased in the leaves of *NtMYB308^CR^* plants, whereas it decreased in NtMYB308OX plants compared to wild-type plants **(Figure 5E)**. The empty vector control displayed GUS staining with almost uniform intensity across all lines. Analysis of relative GUS gene expression further supported these observations, with significantly enhanced expression in *NtMYB308^CR^* and decreased expression in NtMYB308OX compared to wild-type plants **(Figure 5E)**. These findings were consistent with the histochemical staining data, suggesting that as a negative regulator, MYB308 suppresses the activity of *Nt4CL, NtANS*, and *NtCAD*, consequently impacting the expression of the GUS reporter gene in transgenic lines. Overall, these results strongly indicate that the NtMYB308 transcription factor modulates the expression of the GUS reporter gene under the control of Nt4CLpro, NtCADpro, and NtANSpro by regulating their promoter activities.

### NtMYB308 provide tolerance against fungal pathogen *Alterneria solani*

Phenolic compounds, including lignin and other polymers, accumulate in response to both biotic and abiotic stresses, serving as a defense mechanism against pathogen colonization (Kumar et al., 2020; Al-Khayri et al., 2023; Gautam et al., 2023; Caño-Delgado et al., 2003; Tronchet et al., 2010; Sattler and Funnell-Harris, 2013; Buendgen et al., 1990; Bonello et al., 2003). The expression of phenylpropanoid pathway genes and corresponding enzymes is significantly induced during biotic stress, leading to enhanced wall lignification (Kliebenstein et al., 2002; Bhuiyan et al., 2007; Zhao et al., 2009). These insights prompted us to explore the potential role of NtMYB308 in pathogen resistance. To investigate this, leaves of NtMYB308OX, *NtMYB308^CR^*, and wild-type plants were inoculated with conidia of *Alternaria solani* to assess their response to the pathogen. The severity of infection was highest in NtMYB308OX and lowest in *NtMYB308^CR^* plants **(Figure 6A)**. Biotic stress triggers the production of reactive oxygen species (ROS), leading to cell death. Notably, higher staining of NBT and DAB, indicative of ROS accumulation, was observed in NtMYB308OX plants compared to *NtMYB308^CR^* and wild-type plants. **(Figure 6B)**. Additionally, the colony-forming unit (CFU) count per milliliter of culture was measured, revealing higher counts in NtMYB308OX plants **(Figure 6C)**. The reduced lignin content in transgenic plants due to MYB308 overexpression renders them more susceptible to pathogen attacks, as MYB308 acts as a lignin repressor. Conversely, in *NtMYB308^CR^*plants, the loss-of-function of the lignin/phenylpropanoid repressor leads to elevated phenolic content compared to wild-type, providing enhanced resistance against *A. solani*.

**Figure 6.**
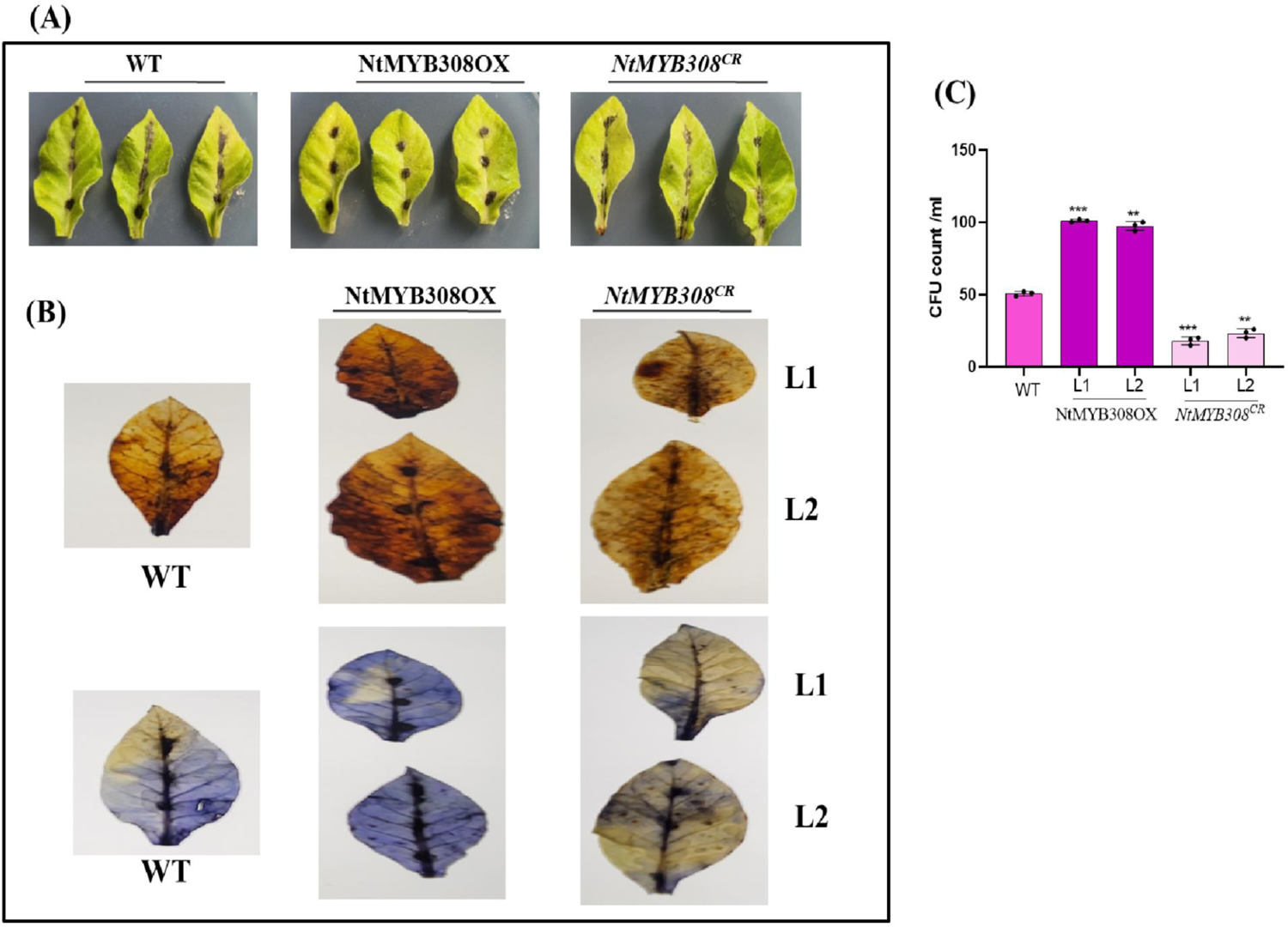
Modulated fungal response in WT, NtMYB308OX and *NtMYB308^CR^* plants. **A**, Representative phenotype of leaves of WT, NtMYB308OX and *NtMYB308^CR^* lines after 5 days of infection with conidia’s of *Alterneia solani* which shows severity of infection in mutant lines. **B,** Detection of Superoxide (O^2-^) by NBT staining and hydrogen peroxide by DAB staining in infected leaves of WT, NtMYB308OX and *NtMYB308^CR^* lines subjected to infection of *A. solani*for 5 days. **C,** CFU (Colony Forming Unit) count in leaves of WT, NtMYB308OX and *NtMYB308^CR^* lines of *Nicotiana tabacum* after 5 days of *A. solani* infection. The experiment was repeated three times independently with similar results. The statistical analysis was performed using two-tailed Student’s t-tests. Data are the means±SE of six biological (n=3) and three technical replicates. The error bars represent standard deviations. The asterisks indicate significant differences: *P<0.1; **P<0.01; ***P< 0.001.

## Discussion

This study identified and characterized three isoforms of R2R3 MYB transcription factors from *Nicotiana tabacum*, affirming their roles as negative regulators of anthocyanin and lignin biosynthesis. R2R3-MYBs exhibit versatility in their function, acting as either activators or repressors of the phenylpropanoid pathway through various mechanisms. These mechanisms include direct binding of transcription factors to the promoters of structural genes or interaction with bHLH and WD40 proteins to form the MBW complex (Naik et al., 2022). Through a series of experiments, we demonstrated that NtMYB308 directly binds to the promoters of *NtANS* and *NtDFR*, inhibiting their expression and subsequently reducing anthocyanin accumulation. Furthermore, we established its direct interaction with the promoters of *Nt4CL* and *NtCAD*, leading to the inhibition of lignin pathway. Overexpression of NtMYB308 in Nicotiana plants resulted in decreased levels of anthocyanin in leaves and flowers, as well as reduced lignin content in the stems of mature plants. Conversely, *NtMYB308^CR^* plants exhibited contrasting outcomes. Additionally, transient silencing of all three isoforms through VIGS yielded similar results to those observed in mutant plants generated via CRISPR/Cas9-based genome editing. These findings underscore the crucial role of NtMYB308 in regulating anthocyanin and lignin biosynthesis pathways, offering valuable insights into the molecular mechanisms underlying plant pigment and lignin production.

A phylogenetic analysis revealed the close relationship between NtMYB308 and its isoforms with AtMYB4-like R2R3 MYBs **(Figure 1B)**, indicating their potential to directly bind to promoters and inhibit anthocyanin and lignin biosynthesis pathways (Yan et al., 2021). Sequence alignment studies further revealed a conserved N-terminal R2R3 domain encompassing a bHLH-binding domain, alongside a highly diverse C-terminal domain shared among all three isoforms, akin to other MYB repressors **(Supplemental Figure 1A)**. Expression analysis of the three identified genes indicated their highest expression levels in leaves, contrary to the conventional expectation of high anthocyanin and lignin accumulation in flowers and stems, respectively **(Figure 1C)**. Interestingly, similar expression patterns were observed in related MYB repressors such as VvMYB4-like, VvMYBC2-L3, PtrMYB182, GbMYBF2, and PhMYB27 (Albert et al., 2011; Xu et al., 2014; Cavallini et al., 2015; Yoshida et al., 2015; Pérez-Díaz et al., 2016), where transcripts were not highly abundant in tissues or developmental stages known for high anthocyanin content. On contrary, studies with *FaMYB1*, *PtrMYB57*, and *Tr-MYB133* showed expression of MYB repressors was highest in tissues with high accumulation of anthocyanin (Aharoni et al., 2011; Albert et al., 2011; Wan et al., 2017). These findings shed light on the intricate regulatory mechanisms underlying anthocyanin and lignin biosynthesis, suggesting an important role for these MYB transcription factors in orchestrating these processes.

Our analysis indicates enhanced expression of anthocyanin and lignin biosynthesis genes in *NtMYB308^CR^* plants **(Figure 3B)**. Conversely, expression of these genes decreased in NtMYB308OX lines compared to WT plants. These results and findings suggest that the modulation of MYB repressor levels affects both early and late flavonoid biosynthesis genes. Similar observations were noted in poplar MYB182OX plants (Yoshida et al., 2015), whereas contrasting results were observed with the overexpression of strawberry FaMYB1 transcription factor (Aharoni et al., 2001), which left the early part of the flavonoid biosynthetic pathway unaffected. Down-regulation of lignin pathway genes leads to negative growth impacts, likely due to metabolic spillover (Besseau et al., 2007; Li et al., 2010; Gallego-Giraldo et al., 2011). These earlier findings prompted us to investigate plant phenotype. Nicotiana NtMYB308OX plants exhibited phenotypic alterations, with significantly reduced size and delayed flowering compared to WT plants **(Figure 3C-F and Supplemental Figure S3C)**. Similar phenotypic outcomes were observed in AmMYB308, AmMYB330, ZmMYB31, PvMYB4-OX, and AtMYB4 overexpression plants (Tamagnone et al., 1998; Fornale et al., 2010; Shen et al., 2012). Reduction in growth of transgenic plants might be the result of challenges in producing mechanical and vascular tissues. Mutants with reduced lignin content, such as Arabidopsis ref8 mutants, also exhibit similar phenotypes (Franke et al., 2002).

Cyanidin, delphinidin, malvidin, petunidin, and pelargonidin are most common anthocyanidins used in various studies (Mattioli et al., 2020; Bloor et al., 1998; Albert et al., 2009). The flower color of overexpression lines exhibited reduced pigmentation, while it was comparatively higher in mutant plants compared to the wild-type **(Figure 4A-B and Supplemental Figure S4A)**. In addition to the flowers, both total anthocyanin and specific anthocyanins were lower in leaves of NtMYB308OX plants and higher in *NtMYB308^CR^* plants compared to wild-type plants **(Figure 4C and Supplemental Figure S4B)**, thus supporting the hypothesis that NtMYB308 acts as an anthocyanin repressor. Similar findings were observed in poplar plant, where MYB182OX lines showed reduced anthocyanin pigmentation in the leaves of transgenic plants compared to wild-type (Yoshida et al., 2015). Clear phenotypic alterations were also evident in petal pigmentation of transgenic plants compared to the control in petunias (MYB27; Albert et al., 2014) and strawberries (FaMYB1; Aharoni et al., 2001). The deficiency in phenolic compounds in plants leads to various physiological changes, such as a reduction in lignin production (Tamagnone et al., 1998). Consequently, there is an interrelation between the reduction in lignin content and cell wall phenolic esters, prompting us to estimate lignin content. Overexpression of NtMYB308, a phenylpropanoid and lignin repressor, resulted in a significant reduction in sinapic acid and syringic acid, whereas they accumulated in high amounts in mutant plants compared to the wild-type, as the mutation led to the non-functional MYB308 **(Figure 4D)**. Similar results were obtained in switchgrass lignin repressor PvMYB4 (Shen et al., 2012) and ZmMYB42, a lignin repressor found in maize (Sonbol et al., 2009), where the content of both syringic acid and sinapate esters were highly reduced in transgenic plants compared to the control. Our analysis also indicated a decrease in lignification within the interfascicular and vascular tissues in the overexpression lines compared to the wild-type. The results concerning total lignin content, vascular bundle length, and lignin staining experiments strongly suggest the role of NtMYB308 as a lignin repressor, as its overexpression resulted in reduced lignin content in transgenic plants. Our findings align with previous studies showing that EgMYB2 acts as a lignin activator when overexpressed (Goicoechea et al., 2005), as higher staining was observed in transformed plants, indicating enhanced lignin content in the xylem, possibly contributing to increased cell wall thickness. In contrast, ZmMYB31, similar to NtMYB308, acts as a lignin repressor; when transformed, the overall lignin content was significantly reduced, as evidenced by the low intensity of staining in transgenic plants (Fornale et al., 2010), reflecting decreased cell wall thickness and altered vascular anatomy. The lignin deposition was highest in mutant and lowest in OX lines when compared to control **(Figure 4F)**. This validates the role of NtMYB308 as lignin repressor. Similar studies of lignin deposition and visualization was done with basic fuschin to investigate the extent of root lignifications by Khandal et al., 2020 and Gaddam et al., 2024.

Both *in silico* analysis and EMSA results revealed the presence of MBSIIG and AC elements upstream of the transcription start site in the promoters of *Nt4CL* and *NtCAD* for lignin biosynthesis, as well as *NtANS* and *NtDFR* for anthocyanin biosynthesis, suggesting potential interactions with NtMYB308. Similar findings have been reported in previous studies, where EgMYB2 was found to regulate the lignin pathway by binding to *EgCAD* and *EgCCR* promoters (Goicoechea et al., 2005). Additionally, Jia et al. (2018) demonstrated the binding of CsMYB308 to *Cs4CL* through transactivation assays. In apple, MdMYB16 and MdMYB6 were shown to regulate anthocyanin metabolism by binding to MBS elements in the *MdANS* promoter, as confirmed by EMSA and Y1H experiments (Xu et al., 2017; Xu et al., 2020). To validate the protein/DNA interaction results and investigate whether NtMYB308 could transcriptionally regulate *Nt4CL*, *NtCAD*, and *NtANS* genes *in vivo*, transactivation assays using the GUS reporter system were conducted in young tobacco leaves. Reduced GUS staining was observed in NtMYB308OX plants infiltrated with *Nt4CL, NtCAD,* and *NtANS* promoters, whereas comparatively higher activity was detected in *NtMYB308^CR^* plants **(Figure 5E)**. These findings parallel previous studies where EgMYB1, acting as a repressor, reduced the activity of *EgCCR* and *EgCAD* promoters (Legay et al., 2007).

Lignin, pectin, and cellulose play crucial roles in positively regulating cell wall integrity, which in turn is essential for plant innate immunity (Ninkuu et al., 2023; Wan et al., 2021). Previous studies have highlighted the significance of molecular switches for lignin in maintaining plant defense against fungi, with mutations in lignin regulators leading to compromised defense mechanisms and decreased plant yield. The involvement of phenolic polymers and lignin in fungal resistance was investigated by infecting leaves with *A. solani*. The severity of infection was found to be highest in NtMYB308OX lines and least in *NtMYB308^CR^* plants. This discrepancy in susceptibility correlates with lignin content, which was lowest in NtMYB308OX plants due to the repressive action of MYB308 on lignin accumulation, while *NtMYB308^CR^* plants exhibited greater tolerance to the fungus. The increased susceptibility observed in NtMYB308OX plants can be attributed to the reduced lignin content, potentially resulting in weaker cell walls. Similar reports from previous studies have demonstrated the impact of fungal pathogens, such as *Cercospora nicotianae*, on tobacco plants. Down-regulation of PAL, leading to reduced chlorogenic acid levels, resulted in severe infection and extensive lesion development (Miedes et al., 2014; Maher et al., 1994). Conversely, transgenic plants constitutively overexpressing PAL genes exhibited high tolerance towards *C. nicotianae* and *Phytophthora parasitica* pv. nicotianae (Way et al., 2002; Shadle et al., 2003). These findings underscore the critical role of lignin and phenolic compounds in plant defense against fungal pathogens and highlight the potential for genetic manipulation to enhance plant resilience.

In conclusion, we present a regulatory network **(Figure 7)** elucidating the role of NtMYB308 as a repressor protein in anthocyanin and lignin biosynthesis through the development of overexpression lines, CRISPR/Cas9-based genome-edited lines, protein-DNA interaction studies, and agro-infiltration-based transactivation studies. The presence of an EAR motif, along with its sequence similarity to subgroup4 MYB transcription factors, further supports its repressor activity. The proposed model suggests that NtMYB308 binds directly to the AC elements of *NtANS, NtDFR, Nt4CL*, and *NtCAD* promoters to regulate anthocyanin and lignin biosynthesis, respectively. The generation of a non-functional MYB repressor protein through CRISPR-Cas9-based genome editing resulted in enhanced anthocyanin and lignin content in mutant plants, leading to enhanced tolerance to fungal infection.

**Figure 7.**
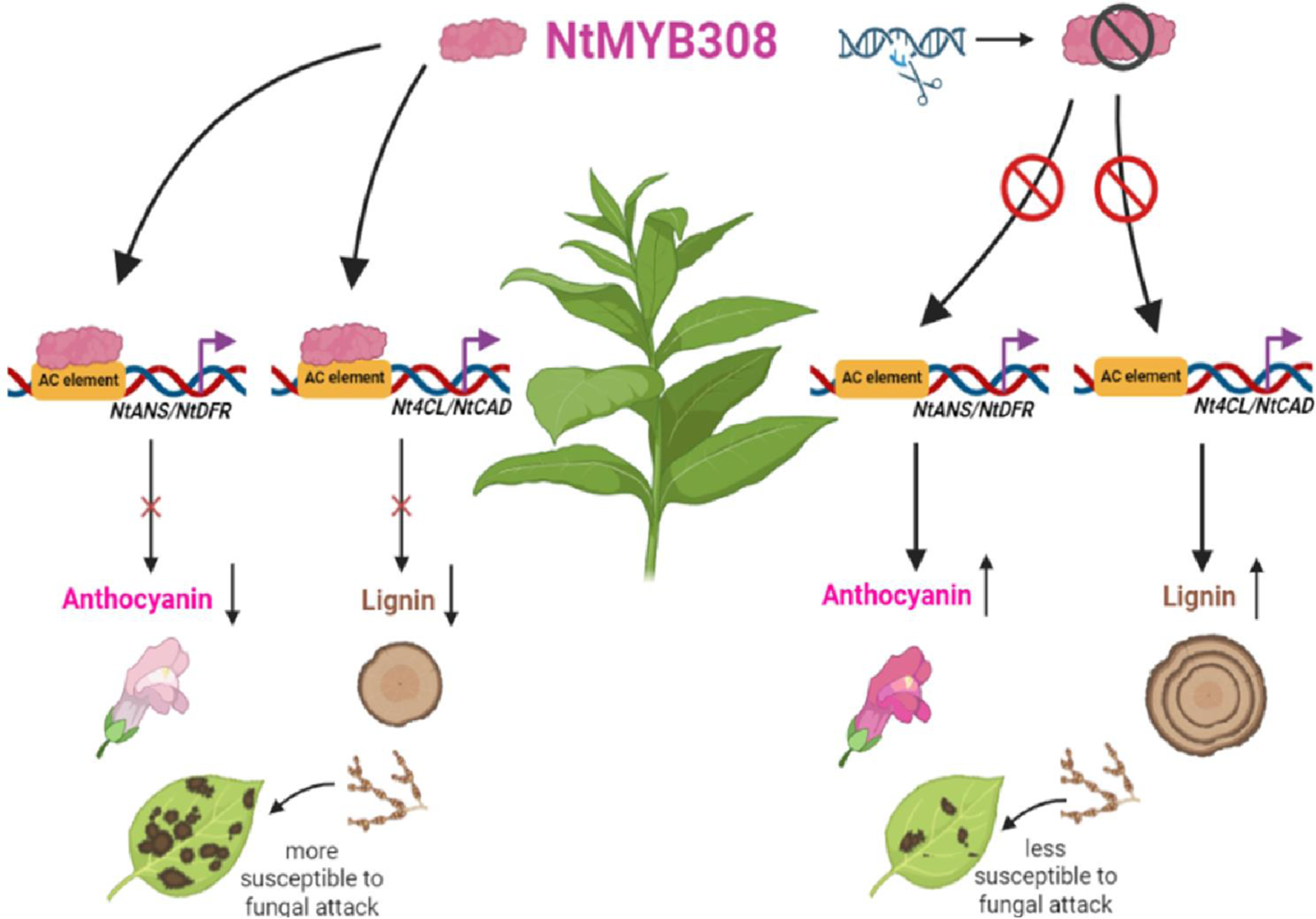
Proposed model showing the regulatory mechanism for anthocyanin and lignin biosynthesis by NtMYB308. NtMYB308 protein of subgroup 4 MYBs contain EAR motif and represses phenylpropanoid/lignin pathway by directly binding to MBSIIG/AC elements in promoters of concerned genes i.e. *NtANS* and *NtDFR*; *Nt4CL* and *NtCAD* promoters and regulate biosynthesis of anthocyanin and lignin respectively. The non-functional MYB repressor protein resulted due to CRISPR-Cas9-based genome-editing inhibits the repression which further leads to enhanced tolerance to fungal infection whereas NtMYB308OX plants are more susceptible to fungal attack.

## Materials and Methods

### Identification of NtMYB308 isoforms and sequence analysis

To identify the ortholog of phenylpropanoid repressors [AtMYB4, AtMYBL2, PtrMYB182 (Poplar), EgMYB1 (Eucalyptus)] in tobacco, BLAST searches were performed on National Center for Biotechnology Information (http://www.ncbi.nlm.nih.gov/BLAST) and amino acid sequence of NtMYB308 and its two isoforms were retrieved and named as NtMYB308-1, NtMYB308-2 and NtMYB308-3. The amino acid sequence was analyzed for N terminal R2R3, C1 and C2 domains and EAR repressor motif, conserved domain database (CCD) was employed to ensure the conserved domains in the identified proteins. The genes retrieved were checked for the presence of the requisite domains using InterProScan (Quevillon et al., 2005) (www.ebi.ac.uk/Tools/pfa/iprscan/). Amino acid sequences of other repressor proteins from different plant species were used to construct the phylogenetic tree using the Neighbor-Joining (NJ) algorithm in MEGA6.0 software (Kumar et al., 2008) and multiple sequence alignment of the aminoacid sequences of R2R3-MYBs was done by ClustalW.

### Plant material, growth conditions and treatments

To raise the transgenic plants seeds were surface sterilized using 70% EtOH for 1 min and then dipped in 50% bleach for 4 min and then washed repeatedly with autoclaved Milli-Q water. They were grown for 2-3 days in absence of light, 24°C temperature, and 60% relative humidity on one-half-strength Murashige and Skoog (MS) medium (Sigma) plates for stratification and after that subsequently transferred to culture room with 16h/8h light-dark photoperiods and 150–180 mm m^-2^s ^-1^ light intensity. *Nicotiana tabacum* cv. Petit Havana (NtPH) leaves were utilized to generate the transgenic plants. In a glasshouse, tobacco plants were cultivated at a temperature of 25°C±2°C with light-dark cycles of 16 hours/8 hours to utilize tissues from various developmental stages.

### Plasmid construction and overexpressing/mutant plant development

Using cDNA as a template and gene-specific primers, VIGS gene fragments (about 150-250 bp) were amplified for NtMYB308-1, NtMYB308-2, and NtMYB308-3 genes. To generate the pTRV2:GOI construct, fragments were cloned into the pTRV2 vector. **Supplemental Table S1** provides the sequences of the oligonucleotides utilized for cDNA amplification and construct development. In this study, the TRV2:NtPDS construct has been generated for NtPDS silencing. Each construct was sequenced to ensure error-free cloning using a capillary automated sequencer (ABI 3730 DNA Analyzer) as per the manufacturer’s instructions. The full-length open reading frame of the NtMYB308-1 cDNA, under the control of the CaMV35S promoter in the binary vector pBI121 (Clontech, USA), was transferred into *Agrobacterium tumefaciens* strain GV3101 and used to transform tobacco plants by leaf disc method (Horsch et al., 1985). Several transgenic tobacco lines constitutively expressing NtMYB308 cDNA were selected on the basis of RT-PCR. Seeds were harvested, sterilised, and plated on solid half strength MS medium supplemented with 100 mg/l kanamycin. Antibiotic-resistant plants were shifted to the glass house and grown until maturity.

CRISPR-Cas9 technology was employed to generate mutant plants of NtMYB308-1, NtMYB308-2 and NtMYB308-3. A 20-bp gRNA was chosen from coding region which was common among all the three isoforms. Thus single guide-RNA was used to edit all the three sequences. BsaI restriction site was used to clone gRNAs into the binary vector pHSE401. The neomycin and hygromycin phosphotransferase genes are included in this vector as selection markers, together with a gene encoding Cas9 endonuclease under the dual CaMV35S promoter. Using plasmid-specific forward and vector reverse primers, all constructs were sequenced from both orientations. They were then transfected to GV3101 and transformed into tobacco.

### Agrobacterium infiltration/VIGS assay

The pTRV1, pTRV2 and pTRV2:GOI constructs were transformed into the LBA4404 strain of *A. tumefaciens* for VIGS assay. A single positive colony was used for primary inoculation and primary culture was grown on 28^ο^C for 16h. Primary culture was then used for secondary culture (50 ml) inoculation which was then grown overnight to get OD600 of 0.6. The secondary culture was then centrifuged at 3,000 x g for 10 min at 4^ο^C and cells were resuspended in infiltration medium (10mM MES, 200µM acetosyringone, 10 mM MgCl_2_) so that OD600 reaches 1.3. Cells were then incubated at 28^ο^C for 3 hours. pTRV1 was co-infiltrated with pTRV2 (Empty vector control), pTRV2:NtPDS and pTRV2:GOI (NtMYB308-1, NtMYB308-2 and NtMYB308-3) in a 1:1 ratio on the abaxial surface of leaves in four-leaf-stage Nicotiana plants. After infiltration, plants were given dark treatment overnight and then kept under controlled growth conditions (25±2^ο^C and 16 h /8 h light/dark treatment) in glass house. The leaves which showed typical viral infection symptoms were collected at 30 d.p.i and stored at −80^ο^C for further analysis. For each construct, 15 independent plants were used in infiltration experiment and six independent infiltrated plants per construct were used for further study.

### Gene expression analysis

Total RNA was isolated using a Spectrum Plant Total RNA kit (Sigma Aldrich) following the manufacturer’s instruction. For qRT-PCR analysis, DNA-free RNA (1 µg) was reverse transcribed using a RevertAid H minus first-strand cDNA synthesis Kit (Fermentas) according to the manufacturer’s instructions. qRT-PCR was performed using Fast Syber Green mix (Applied Biosystems) in a Fast 7500 Thermal Cycler instrument (Applied Biosystems). Expression was normalized using Tubulin and analyzed through the comparative CT method (Liva and Schmittgen, 2001). The oligonucleotide primers used to study expression of different genes were designed using the Primer Express 3.0.1 tool (Applied Biosystems) and information is provided in **Supplemental Table S1**

### Quantification of total anthocyanin

To extract anthocyanin, 300 mg of tissue were boiled for three minutes in an extraction solution (propanol: HCl: H2O, 18:1:81), and then the tissue was left to stand at room temperature in the dark for a minimum of two hours. After centrifuging the samples, the supernatants’ absorbance was measured at 650 and 535 nm.

### Extraction and quantitative estimation of anthocyanins

Fresh petal tissues weighing 300mg were finely ground in liquid nitrogen and 1ml extraction buffer (0.5% HCl in methanol:water, 1:1) was added and vortexed. The tube was centrifuged for 10 minutes at 10,000 rpm after being left in the dark for 30 minutes. Supernatant was pipette out into a fresh tube containing 0.2 mL of chloroform. After 30 seconds of vortexing, the tube was centrifuged for 2 minutes at 10,000 rpm. The non-polar compound-containing chloroform phase was removed and this process was repeated. Applying a UV spectrophotometer (UV-2550, Shimadzu, Kyoto, Japan), the upper methanol and water phase was pipetted into a fresh tube for anthocyanin measurement at 530 nm. Next, in a 1.5 mL tube, 50 μL of anthocyanins extract was combined with 950 μL of butanol:HCl (95:5, v/v). After a one-hour boil, the mixture was allowed to cool to ambient temperature before being quickly evaporated in a speed vacuum. For HPLC analysis, 50 μL of methanol containing 0.1% HCl was used to suspend the leftover anthocyanidin residue.0.1% acetic acid (solvent A) and acetonitrile (solvent B) made up the elution solvent system. In order to elute anthocyanidins, the gradient programme consists of various ratios of solvent A to solvent B. 90:10 to 83:17 (0–5 min), 83:17 to 77:23 (5–10 min), 77:23 to 71:29 (10–15 min), 71:29 to 68:32 (15–20 min), 68:32 to 65:35 (20–25 min), 60:35 (25–39 min), 65:35 to 50:50 (39–45 min), 50:50 to 70:30 (45–50 min), and 70:30 to 90:10 (50–55 min) were the sequence of events. Ten minutes were dedicated to column washing afterward. The injection volume was 5 μL, and the flow rate was 1 mL/min. The chromatogram was taken at 530 nm. As positive controls, three standards were injected: cyanidin chloride, delphinidin chloride, malvidin and pelargonidin chloride (Zhu et al., 2018).

### Visualization and extraction of total lignin

The Leica DM2500 microscope70 was used to view the hand-cut sections that had been treated with phloroglucinol (Sigma-Aldrich) for one minute in order to visualise the lignified cells in the stems. For quantification of lignin, (Bruce and West, 1989) was followed. In a nutshell, the 600 mg of samples were pulverised in liquid N_2_ and suspended in 2 ml of ethanol. At 12,000xg, the mixture was centrifuged for 30 minutes at 4 °C. After being dried and resuspended in 5 ml of 2N HCl and 0.5 ml of thioglycolic acid, the pellet was incubated for 8 hours at 95°C before being cooled to room temperature. After centrifuging the suspension for 30 minutes at 12,000xg, double-distilled water was used to wash the pellet. The pellet that resulted from another centrifugation at 12,000xg and 4°C for 5 minutes was suspended in 5 ml of 1N NaOH and gently stirred at 25 °C for eighteen hours. One ml of HCl was added to the supernatant following a 30-minute centrifugation at 12,000xg, 4°C. The mixture was then left to precipitate at 4°C for an entire night. The pellet was redissolved in 3 ml of 1N NaOH after centrifugation at 12,000xg, 4°C for 30 minutes, and the amount of lignin was measured using absorbance at A280.

### Extraction of phenolic content through HPLC

Xylem tissue (300 mg) was acquired by scraping the bark-free stems of 45 days old Nicotiana plants grown in soilrite and it was subsequently homogenized in liquid nitrogen, and an initial extraction was performed with 5 ml of methanol and then with cyclohexane/water containing 0.1% trifluoroacetic acid (1:1, v/v). For the analysis of phenolic compounds, reversed-phase High-Performance Liquid Chromatography (HPLC) was employed (Meyermans et al., 2000). The separation was achieved through a gradient elution with methanol-acetonitrile (25:75, v/v) acidified with 0.1% trifluoroacetic acid (solvent B) in 0.1% aqueous trifluoroacetic acid (solvent A). The gradient elution conditions were as follows: 0 min/0% B, 25 min/60% B, 27 min/100% B. The HPLC conditions included a flow rate of 1.5 ml/min, a temperature of 40 °C, and an injection loop of 20 ml. All solvents used were of HPLC grade purity.

### Expression and purification of NtMYB308 protein

The entire open reading frame of NtMYB308-1 was amplified and cloned into the pET-28b(+) vector (Novagen, Germany). This construct was introduced into *E. coli* BL21 (DE3) pLysS (Invitrogen, USA) for prokaryotic expression, induced with IPTG. The recombinant 6X-His-tagged NtMYB308 protein was purified using a Ni-NTA column (Nucleopore, India), and the eluted protein concentration was determined using a Bradford assay.

### Electrophoretic Mobility Shift Assay

The labeling of probes with digoxigenin was performed using the 2nd generation DIG Gel Shift EMSA kit (Roche, USA), following the instructions provided by the manufacturer. Then labelled probes were incubated for 30-minute at 21°C in a binding buffer comprising 100 mM HEPES (pH 7.6), 5 mM EDTA, 50 mM (NH_4_)_2_SO_4_, 5 mM DTT, 1% (w/v) Tween 20, and 150 mM KCl, with and without the presence of the recombinant protein. To evaluate specific binding, reactions were set for increasing concentrations of unlabeled probes. Afterward, the binding reaction was separated on a 6% polyacrylamide gel in 0.5X TBE (pH 8.0) buffer and semi-dry transferred onto a positively charged nylon membrane using the Transblot system from BIO-RAD, USA. UV cross-linking was performed on the transferred material. Finally, the membrane was incubated with CSPD chemiluminescent solution and was exposed to X-ray blue film (Retina, India) for subsequent analysis.

### Transient expression in tobacco

The promoter constructs viz ProNt4CL::GUS, ProNtANS::GUS, ProNtCAD::GUS, and EV (empty vector positive control which includes β-glucuronidase gene under the control of the CaMV35S promoter), were transformed into *Agrobacterium tumefaciens* (strain GV3101) and used for agro-infiltration. Agrobacteria strains were grown at 28^ο^C and resuspended at OD600=0.6 in infiltration medium (10 mM MgCl_2_, 10 mM MES and 200 µM acetosyringone) and incubated for 3 h at room temperature (Gani et al., 2022). The induced Agrobacterium cultures were injected into the near fully expanded leaves of Nicotiana plantlets (control, NtMYB308OX and *NtMYB308^CR^*) grown for 4-5 weeks in controlled conditions. Prior to the histochemical GUS assays, the infiltrated plants were kept in a growth chamber for three to four days. The histochemical GUS staining procedure was carried out according to Jefferson et al. (1989).

### Statistical analysis

The statistical tests and n numbers, including sample sizes or biological replications, are described in the figure legends. All the statistical analyses were performed using two-tailed Student’s t-tests using GraphPad Prism version 9.0 software. All the experiments were repeated at least three times independently, with similar results.

### Gene accession numbers

NtMYB308-1 (XM_016613474), NtMYB308-2 (XM_016632216), NtMYB308-3 (XM_016645944)

## Supplemental data

**Supplemental Figure S1.** Nucleotide and protein sequence alignment of three isoforms of NtMYB308 involved in anthocyanin and lignin regulation

**Supplemental Figure S2.** Phenotypic changes and modulation in metabolite content in TRV mediated silenced lines of three homologs of NtMYB308

**Supplemental Figure S3.** Schematic representation of genome editing in *NtMYB308^CR^* edited WT tobacco plants and down-regulation of three isoforms in respective CRISPR/Cas9-based genome-edited lines.

**Supplemental Figure S4.** NtMYB308 modulates total anthocyanin content in flower and leaves as well as total lignin content in stem of genome-edited and transgenic plants.

**Supplemental Figure S5.** Recombinant protein expression and purification of NtMYB308 and NtMYB308 physically binds to Nt4CL, NtCAD and NtANS, NtDFR promoters.

**Supplemental Figure S6.** *In silico* analysis of *Nt4CL, NtCAD, NtANS* promoters

## Data availability

All data generated or analyzed during this study are included in this published article (and its supplementary information files).

## Acknowledgment

P.K.T. acknowledges CSIR New Delhi, for financial support in the form of projects on Plant Genome Editing. P.K.T. also acknowledges Science and Engineering Research Board, New Delhi for JC Bose National Fellowship (JCB/2021/000036). S.D. and P.K.P. acknowledge DBT, New Delhi for Senior and Junior Research Fellowship respectively. Authors also acknowledge Dr. Manju Singh. CSIR-CIMAP and Dr. Suchi Srivastava from CSIR-NBRI for help in metabolite profiling and fungal bioassay, respectively.

